# Primary Cilia Loss in Striatal Neurons Disrupts Synaptic Connectivity and Excitatory Transmission and Drives Metabolomic Remodeling

**DOI:** 10.64898/2026.07.30.741876

**Authors:** Kiki Chen, Archana Proddutur, Dana Shevachman, Xiangrong Feng, Sammy Alhassen, Rouda Vakil Monfared, Wedad Alhassen, Travis Dabbous, Joshua Lee, Surya Nauli, Kevin Beier, Gyorgy Lur, Amal Alachkar

**Affiliations:** Department of Pharmaceutical Sciences, School of Pharmacy, University of California-Irvine, CA 92697, USA; Department of Neurobiology and Behavior, University of California, Irvine, CA, USA 92697-4560; Department of Biomedical and Pharmaceutical Sciences, School of Pharmacy, Health Science Campus, Chapman University, Irvine, USA; Department of Physiology and Biophysics, School of medicine, University of California, Irvine, USA; Department of Biomedical Engineering, University of California, Irvine, CA, USA 92697-4560; UC Irvine Center for the Neurobiology of Learning and Memory, University of California Irvine, Irvine, California, CA 92697, USA; UC Irvine Artificial Intelligence in Science Institute, University of California Irvine, Irvine, California, CA 92697, USA

**Keywords:** primary cilia, IFT88, striatum, monosynaptic rabies tracing, electrophysiology, metabolomics, basal ganglia, synaptic connectivity, NMDA receptor, diacylglycerol

## Abstract

Disruption of striatal circuits is a central feature of many neurological and psychiatric disorders, yet the mechanisms that maintain afferent connectivity and synaptic function in striatal neurons remain incompletely defined. Primary cilia are signaling organelles present on almost all striatal medium spiny neurons that are enriched in neuromodulatory receptors, suggesting a role in coordinating striatal neuronal communication and biochemical state. Here, we show that conditional ablation of primary cilia from striatal neurons by AAV-Cre-mediated deletion of *Ift88* disrupts the afferent connectivity, synaptic function, and chemical signature of the striatum. Monosynaptic rabies tracing revealed an approximately threefold reduction in brain-wide input convergence onto striatal neurons. Whole-cell recordings showed reduced miniature excitatory postsynaptic current amplitude and frequency together with a reduced NMDA:AMPA ratio, consistent with weakened glutamatergic synaptic transmission. Untargeted metabolomics revealed broad remodeling of the striatal chemical profile, predominantly toward decreased measured levels, with lipid-associated pathways most affected alongside reductions in polyamines, glutamate-related metabolites, and neuromodulatory, particularly excitatory, signaling molecules. By contrast, the cortex, which was not targeted by the AAV injection, showed fewer and directionally opposite molecular changes. These findings identify cilia as essential regulators of the structural, synaptic, and molecular integrity required for normal striatal circuit function.

## INTRODUCTION

Disruption of striatal circuits is a central feature of many neurological and psychiatric disorders, yet how these circuits maintain their connectivity and function in the adult brain remains incompletely understood. As the principal input nucleus of the basal ganglia, the striatum integrates widespread glutamatergic afferents from the cortex and thalamus, together with dopaminergic and other neuromodulatory inputs, to regulate action selection, reward learning, habit formation, and behavioral flexibility [1–5].

Striatal medium spiny neurons (MSNs) are the main site of integration, and convey information to the output nuclei of the basal ganglia [2, 6]. Nearly all MSNs possess a primary cilium, a solitary, non-motile, microtubule-based organelle that functions as a specialized signaling compartment [7, 8]. Once regarded as vestigial, they are now recognized as specialized signaling compartments that concentrate receptors and effectors of multiple pathways, including Hedgehog, Wnt, and G protein-coupled receptor (GPCR) signaling, allowing spatially-restricted integration of extracellular cues [9–13]. This compartmentalization enables cilia to act as localized signaling platforms enriched in GPCRs and second-messenger machinery that shape neuronal state, synaptic input, and circuit organization [14, 15]. In MSNs, these cilia are enriched in neuromodulatory receptors, including dopamine receptors D1 (DRD1) and D2 (DRD2), somatostatin receptor 3 (SSTR3), melanocortin receptor 4 (MC4R), melanin-concentrating hormone receptor 1 (MCHR1), and Smoothened (Smo) [10, 14, 16–19], positioning the MSN cilium as a hub through which neurochemical cues can be coordinated with afferent input.

Beyond development, neuronal cilia also regulate mature neuronal structure and function, influencing dendritic morphology, synaptic integration, and behavior, with forebrain-wide cilia ablation disrupting cognitive, affective, and sensorimotor processes [20–22]. We previously showed that selective ablation of striatal cilia in adult mice impairs motor-skill learning, sensorimotor gating, spatial working memory, and short-term social memory, increases repetitive grooming, and alters neuronal activity in the dorsal striatum and in cortical regions that project to it [23]. These findings established that striatal cilia are necessary for multiple dorsal-striatum-dependent behaviors and for normal corticostriatal activity. However, how striatal cilia regulate the cellular, synaptic, and chemical organization that underlies these functions remains unknown.

To address this gap, we selectively removed primary cilia from dorsal striatal neurons via AAV-Cre-mediated deletion of Ift88 [24], and combined monosynaptic rabies tracing, whole-cell electrophysiology, and untargeted metabolomics to examine afferent connectivity, excitatory synaptic transmission, and molecular state. We find that loss of striatal cilia leads to a broad reduction in presynaptic inputs, weakens excitatory synaptic drive onto medium spiny neurons, and induces pronounced metabolomic remodeling. These results demonstrate that primary cilia are essential for maintaining the structural, synaptic, and molecular integrity required for proper excitatory integration in the adult striatum.

## RESULTS

### Efficient cilia ablation in striatal neurons without affecting starter cell numbers

To ablate primary cilia selectively in striatal neurons, Ift88*fl/fl* mice received bilateral stereotaxic co-injection of AAV9- CaMKII-Cre together with helper viruses (TVA/RG) into the dorsal striatum, and WT controls (Ctrl) received identical injections (Fig. 1A,B). Expression of TVA enables selective infection of TVA-expressing cells using an EnvA-pseudotyped rabies virus (RABV), and RG enables trans-complementation and monosynaptic spread to input cells [25]. Cilia ablation was confirmed by quantification of ADCY3-positive cilia in TVA-expressing neurons at the injection site (Fig. 1C). cKO mice showed a ∼75% reduction in cilia relative to controls (Ctrl: 100.0 ± 5.0%; cKO: 24.9 ± 6.8%; p = 0.0009; Fig. 1C,D), confirming efficient and selective cilia ablation in transduced striatal neurons.

**Figure 1.**
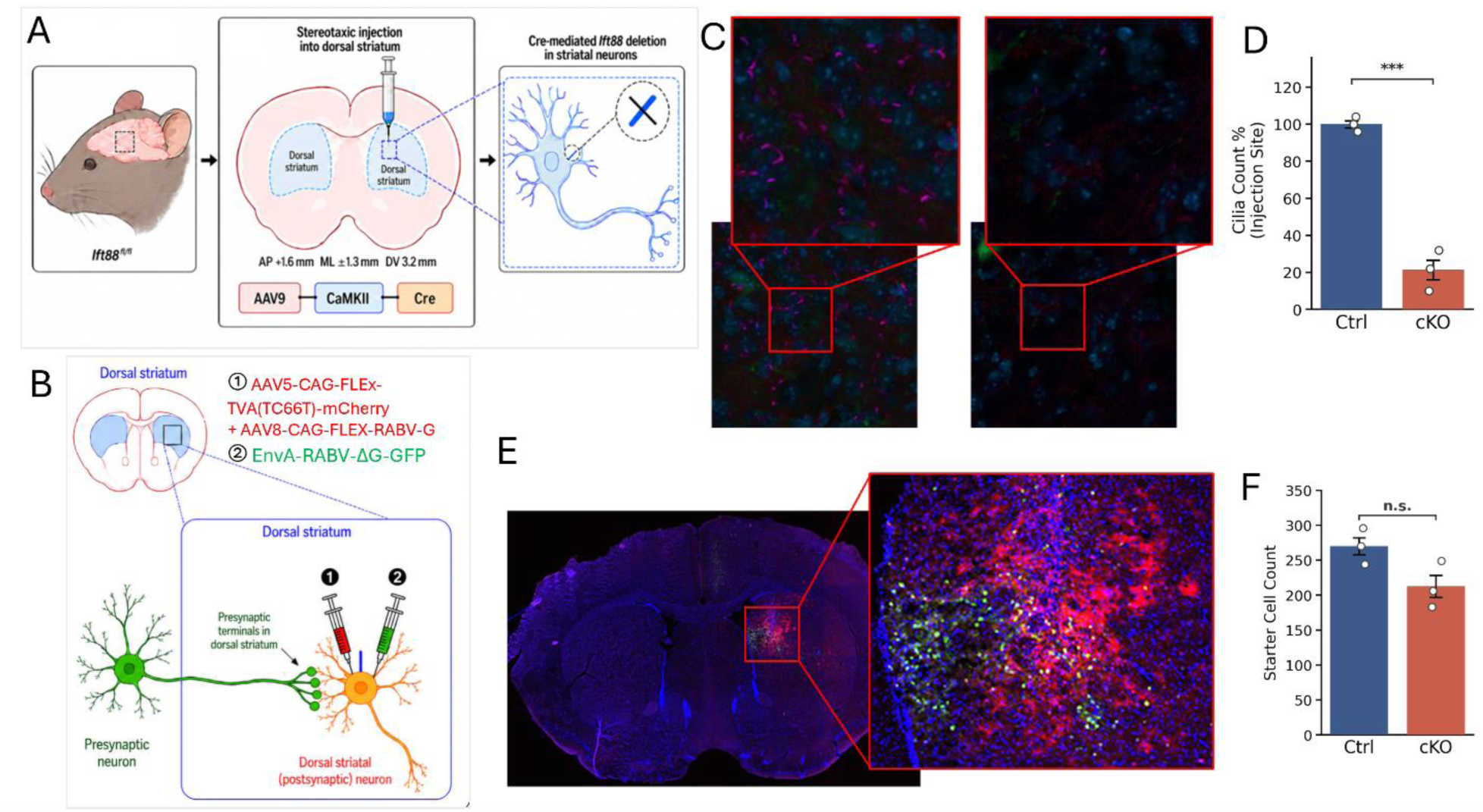
Selective ablation of primary cilia in striatal neurons does not affect starter cell generation. **(A)** Schematic of the experimental approach. Ift88^fl/fl^ mice received bilateral stereotaxic injection of AAV9-CaMKII-Cre into the dorsal striatum to induce Cre-mediated deletion of Ift88 in striatal neurons, hereafter referred to as conditional knockout (cKO). Wild-type controls (Ctrl) received the same viral injections without Cre-dependent Ift88 deletion. **(B)** Schematic of the monosynaptic rabies tracing strategy. Helper viruses encoding TVA and rabies glycoprotein (RG) were co-injected into the dorsal striatum to define starter cells, followed by EnvA-pseudotyped, glycoprotein-deleted rabies virus to label monosynaptic presynaptic inputs. **(C)** Representative confocal images showing ADCY3-immunolabeled primary cilia in TVA-expressing neurons at the injection site in Ctrl and cKO mice. Insets show higher-magnification views. Scale bars as indicated. **(D)** Quantification of ADCY3-positive cilia at the injection site showing reduced cilia counts in cKO mice compared with Ctrl mice. Data are mean ± SEM; n = 3 mice per group; unpaired t-test, t = 8.844, **p = 0.0009. **(E)** Representative confocal images showing TVA-mCherry^+^/GFP^+^ double-positive starter neurons at the injection site. Insets show higher-magnification views of starter-cell populations. **(F)** Quantification of starter-cell counts showing no significant difference between Ctrl and cKO mice. Data are mean ± SEM; n = 3 mice per group; unpaired t-test, n.s.: non-significant, p > 0.05.

To determine whether cilia loss altered viral access or starter cell generation, we quantified TVA-mCherry⁺/GFP⁺ double- positive neurons at the injection site. Starter cell counts were comparable between Ctrl and cKO mice (Ctrl: 270 ± 20; cKO: 213 ± 19; n.s.; Fig. 1E,F), confirming equivalent AAV transduction efficiency across groups.

### Loss of striatal cilia causes a marked reduction in brain-wide monosynaptic input convergence with reorganization of afferent input distribution

To assess whether loss of striatal cilia alters overall monosynaptic input convergence, we quantified total input cell counts and normalized input strength by starter cell number (Fig. 2A-B). Total input cell counts were markedly reduced in *Ift88* cKO mice relative to controls (Ctrl: 3267 ± 279; cKO: 854 ± 135; p = 0.0019; Fig. 2C). Similarly, the input/starter ratio was significantly reduced in cKO mice (Ctrl: 12.30 ± 0.42; cKO: 4.29 ± 0.91; p = 0.0013; Fig. 2D), representing an approximately 3-fold decrease. These findings indicate that loss of striatal cilia is associated with a robust reduction in overall monosynaptic input convergence in MSNs.

**Figure 2.**
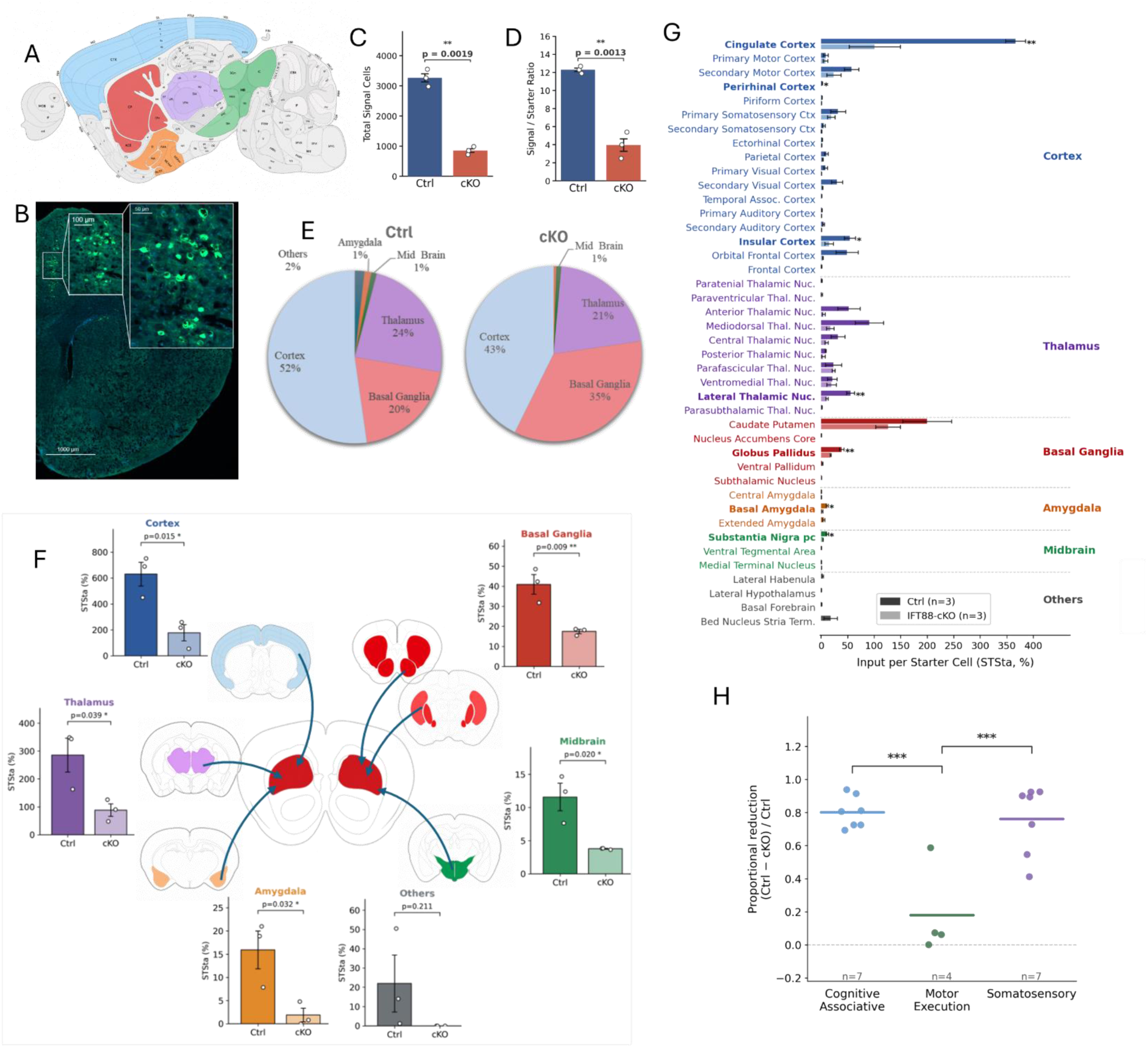
Loss of striatal cilia reduces brain-wide monosynaptic input convergence and selectively alters afferent circuit architecture. **(A)** Schematic illustration of a representative coronal brain section showing the major afferent region groups (color-coded) that provide monosynaptic input to the dorsal striatum. **(B)** Representative whole-brain coronal fluorescence image showing retrogradely labeled presynaptic input neurons (GFP+, green) in an IFT88-cKO mouse. Inset shows a higher-magnification view of labeled neurons at the injection site. Scale bars: 1000 μm (main), 100 μm (inset). **(C)** Total signal cell counts in cKO and control mice (unpaired t-test; p = 0.0019). **(D)** Signal-to-starter ratio (STSta) was significantly reduced in cKO mice (unpaired t-test; p = 0.0013), representing an approximately threefold decrease. **(E)** Pie charts showing the proportional contribution (%) of major brain region groups to the total afferent pool in Ctrl and cKO mice. Proportions are within-animal percentages, shown for description only; no statistical test is applied to the composition. Data are presented as mean ± SEM. n = 3 mice per group. **(F)** Absolute starter-cell-normalized input strength (STSta) across six major brain-region classes; Unpaired two-tailed t-test; *p < 0.05, n.s.: non-significant. **(G)** Region-level analysis of starter-cell-normalized input strength across individual afferent regions. **(H)** Circuit-domain analysis showing proportional reduction of input strength, calculated as (Ctrl − cKO)/Ctrl, across cognitive-associative, motor execution, and somatosensory domains. Regions were assigned to domains as follows — Cognitive-associative (n = 7): Cingulate Cortex, Orbital Frontal Cortex, Insular Cortex, Mediodorsal Thalamic Nucleus, Central Thalamic Nucleus, Anterior Thalamic Nucleus, Lateral Thalamic Nucleus; Motor execution (n = 4): Primary Motor Cortex, Secondary Motor Cortex, Ventromedial Thalamic Nucleus, Parafascicular Thalamic Nucleus; Somatosensory (n = 7): Primary Somatosensory Cortex, Secondary Somatosensory Cortex, Parietal Cortex, Primary Visual Cortex, Secondary Visual Cortex, Secondary Auditory Cortex, Posterior Thalamic Nucleus. Domain differences were analyzed by one-way ANOVA followed by Tukey HSD post hoc test: F(2,15) = 15.741, p = 0.0002; cognitive-associative vs. motor execution, p = 0.0003; motor execution vs. somatosensory, p = 0.0005; cognitive-associative vs. somatosensory, p = 0.917. Data are presented as mean ± SEM; n = 3 mice per group. *p < 0.05, **p < 0.01, ***p < 0.001.

Although total monosynaptic input was markedly reduced in cKO mice, the reduction was not uniform across input sites (Fig. 2E). For example, the proportional contribution of cortical input decreased (∼52% in controls to ∼43% in cKO), whereas the basal ganglia share increased (∼20% to ∼35%; Fig. 2E). Because this compositional analysis includes local intrastriatal inputs within the basal ganglia class, the increased basal ganglia share reflects relative sparing of local striatal connectivity as long-range input declined.

To identify which afferent systems contributed to the reduction in input, retrogradely labelled inputs were grouped into six major brain region classes: cortex, thalamus, basal ganglia (extra-striatal nuclei only), amygdala, midbrain, and others (lateral habenula, lateral hypothalamus, basal forebrain, bed nucleus of stria terminalis). Analysis of absolute starter-cell-normalized input strength (STSta) revealed significant reductions in five of six groups (Fig. 2F, Supplementary Table 1). Cortical input showed the largest absolute decrease (Ctrl: 629.9 ± 91.8 vs. cKO: 177.1 ± 62.4; p = 0.015). Thalamic input was also significantly reduced (Ctrl: 285.3 ± 60.9 vs. 88.3 ± 22.5; p = 0.039), as were inputs from the amygdala (Ctrl: 15.9 ± 4.1 vs. 1.9 ± 1.5; p = 0.032) and Midbrain (Ctrl: 11.6 ± 2.1 vs. 3.8 ± 0.1; p = 0.020, Fig. 2F). Inputs from extra-striatal basal ganglia nuclei were also significantly reduced in cKO mice (Ctrl: 40.9 ± 4.9 vs. 17.4 ± 1.0; p = 0.009), though proportionally less than cortical input (57% vs 72% reduction). The ‘Others’ category showed a non-significant trend towards reduction (Ctrl: 22.0 ± 14.7 vs. 0.0 ± 0.0; p = 0.211).

Region-level analysis identified 7 individual brain areas with significantly reduced input following striatal cilia ablation (Fig. 2G), and no region showed a significant increase. Cortical reductions were most prominent in the cingulate cortex (Ctrl: 366.5 ± 18.8% vs. cKO: 100.9 ± 48.2%; p = 0.007), insular cortex (54.0 ± 10.8% vs. 14.9 ± 8.1%; p = 0.044; Fig. 2G), and perirhinal cortex (p = 0.021). Thalamic input was significantly reduced in the lateral thalamic nucleus (55.2 ± 7.7% vs. 10.7 ± 2.3%; p = 0.005; Fig. 2G). Significant reductions were also detected in the lateral globus pallidus (p = 0.008), basolateral amygdala (p = 0.014), and substantia nigra pars compacta (p = 0.017; Fig. 2G). Together, these findings indicate that the loss of input is distributed across multiple convergent afferent systems rather than restricted to a single pathway.

Analysis by functional circuit domain revealed marked selectivity in the pattern of input loss (Fig. 2H). Cognitive-associative inputs were most severely affected, showing a mean proportional reduction of 80.2 ± 3.6%, with particularly strong losses from the cingulate, orbitofrontal, and insular cortices as well as higher-order thalamic nuclei (mediodorsal, anterior, and lateral). Somatosensory inputs were also strongly reduced (76.1 ± 7.9%), especially from primary and secondary somatosensory cortices and visual areas. In contrast, motor execution inputs were only modestly affected (18.1 ± 13.6%), with relatively preserved contributions from primary and secondary motor cortices and ventromedial thalamus. This differential vulnerability was statistically significant, with motor execution circuits being significantly more spared than both cognitive-associative (p = 0.0003) and somatosensory (p = 0.0005) domains. These findings demonstrate that striatal cilia ablation preferentially disrupts associative and sensory afferents while largely sparing motor execution pathways (Fig. 2H).

### Cilia ablation reduces excitatory synaptic transmission in striatal MSNs

To determine whether cilia ablation in dorsal striatal neurons caused changes in excitatory synaptic drive to MSNs, we first recorded miniature excitatory postsynaptic currents (mEPSCs) from these neurons (Fig 3A-D). Selective ablation of cilia from the dorsal striatum caused a significant decrease in mEPSC amplitude in cKO group compared to control mice (Ctrl: 37.34±2.4 pA; median = 38.19, IQR = 32.4 - 43.66, n = 12cells/ 6mice; cKO: 30.36±3.8 pA, median = 28.8, IQR = 20.97 – 36.13, n = 8 cells/ 3mice, p < 0.05, two sample Kolmogorov-Smirnov test; Fig 3E and G). Additionally, we observed a significant reduction in the frequency of mEPSCs when compared to Ctrl mice (Ctrl: 22.76±2.9 Hz; median = 24.08, IQR = 12.95 - 29.14, n = 12 cells/ 6 mice; cKO: 12.62±2.5 Hz, median = 10.86, IQR = 7.8 - 19.2, n = 8 cells/ 3mice, p<0.05, two sample Kolmogorov-Smirnov test; Fig 3F and H). These findings indicate a decrease in the number of functional synapses and impaired glutamatergic transmission on medium spiny neurons.

**Figure 3.**
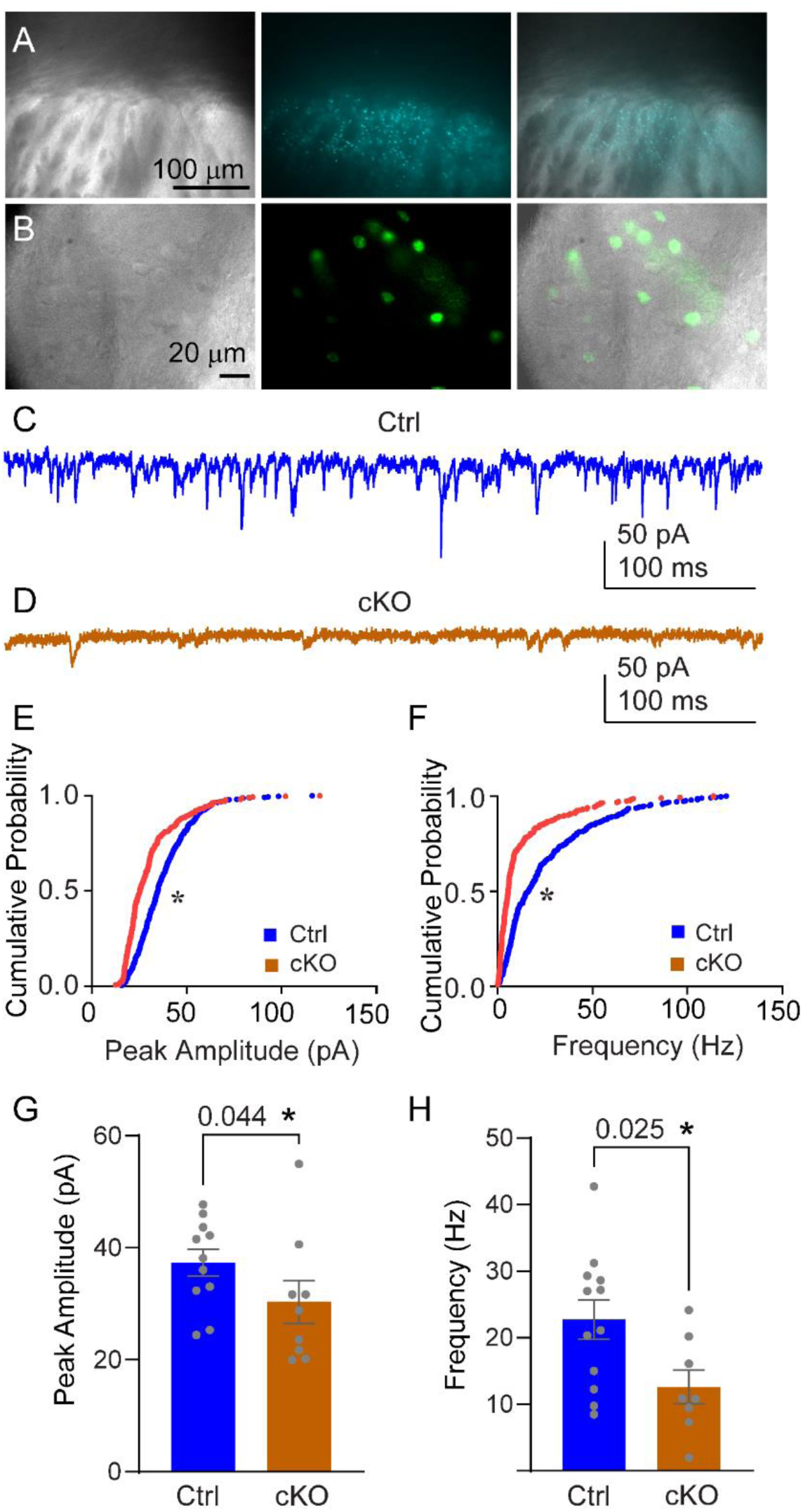
Cilia ablation reduces miniature excitatory postsynaptic current amplitude and frequency in striatal MSNs. **(A)** Visualization of viral expression in acute dorsal striatal slices. Live 400-μm coronal striatal slices were imaged at 10X magnification. Left: IR- DIC images, middle: GFP fluorescence, right: overlay showing the broader striatal injection area and expression pattern within the slice. (**B**) high magnification (60X) images show the cellular morphology used for targeting medium spiny neurons (left). Middle: fluorescence images acquired using the GFP cube. Right: merged DIC and GFP channels illustrate how the recorded neurons were targeted. **(C)** Representative mEPSC traces recorded from striatal neurons in control and **(D)** cKO mice. Scale bars: 50 pA, 100 ms. **(E)** Cumulative probability distribution of mEPSC peak amplitude for Ctrl and cKO cells. *p < 0.05, two-sample Kolmogorov- Smirnov test. **(F)** Cumulative probability distribution of mEPSC inter-event interval; *p < 0.05, two-sample Kolmogorov-Smirnov test. **(G)** Mean mEPSC peak amplitude. **(H)** Mean mEPSC frequency. Data in bar graphs are presented as mean ± SEM with individual data points. n = 12 cells/6 mice Ctrl, n = 8 cells/3 mice cKO. Unpaired t-test.

### Cilia ablation alters the NMDA receptor activation

To further characterize the impact of cilia ablation on excitatory synaptic transmission in dorsal striatal MSNs, we recorded evoked AMPA- and NMDA receptor-mediated currents (Fig. 4A-C). The NMDA:AMPA ratio reflects the relative contribution of these receptor types to glutamatergic transmission at excitatory synapses. We observed a significant decrease in NMDA:AMPA ratio in the cKO group compared to control mice (Ctrl: 1.67±0.23; cKO: 0.77±0.22; p = 0.012, unpaired t-test, n = 9 cells from 4 mice per group; Fig. 4D, Supplementary Table 2). We did not observe changes in paired pulse ratio for either glutamatergic current (Fig. 4E and F), indicating likely postsynaptic mechanisms. Peak amplitude of AMPA receptor mediated responses did not show a statistically significant difference (p = 0.13, Fig. 4G) while the amplitude of NMDA receptor responses showed a trend towards reduced activation (p = 0.083, Fig. 4J) in cKO mice. Response kinetics of AMPA receptor mediated currents also remained unchanged (Fig 4H, I). In contrast, NMDA currents showed a trend towards slower rise (p = 0.082, Fig 4K) and markedly slowed decay (Ctrl: 71.34 ± 7.92 ms; cKO: 102.35 ± 11.30 ms, p = 0.039, Fig. 4L) kinetics with cilia ablation. These findings indicate that cilia ablation selectively impacts glutamatergic transmission via NMDA receptors.

**Figure 4.**
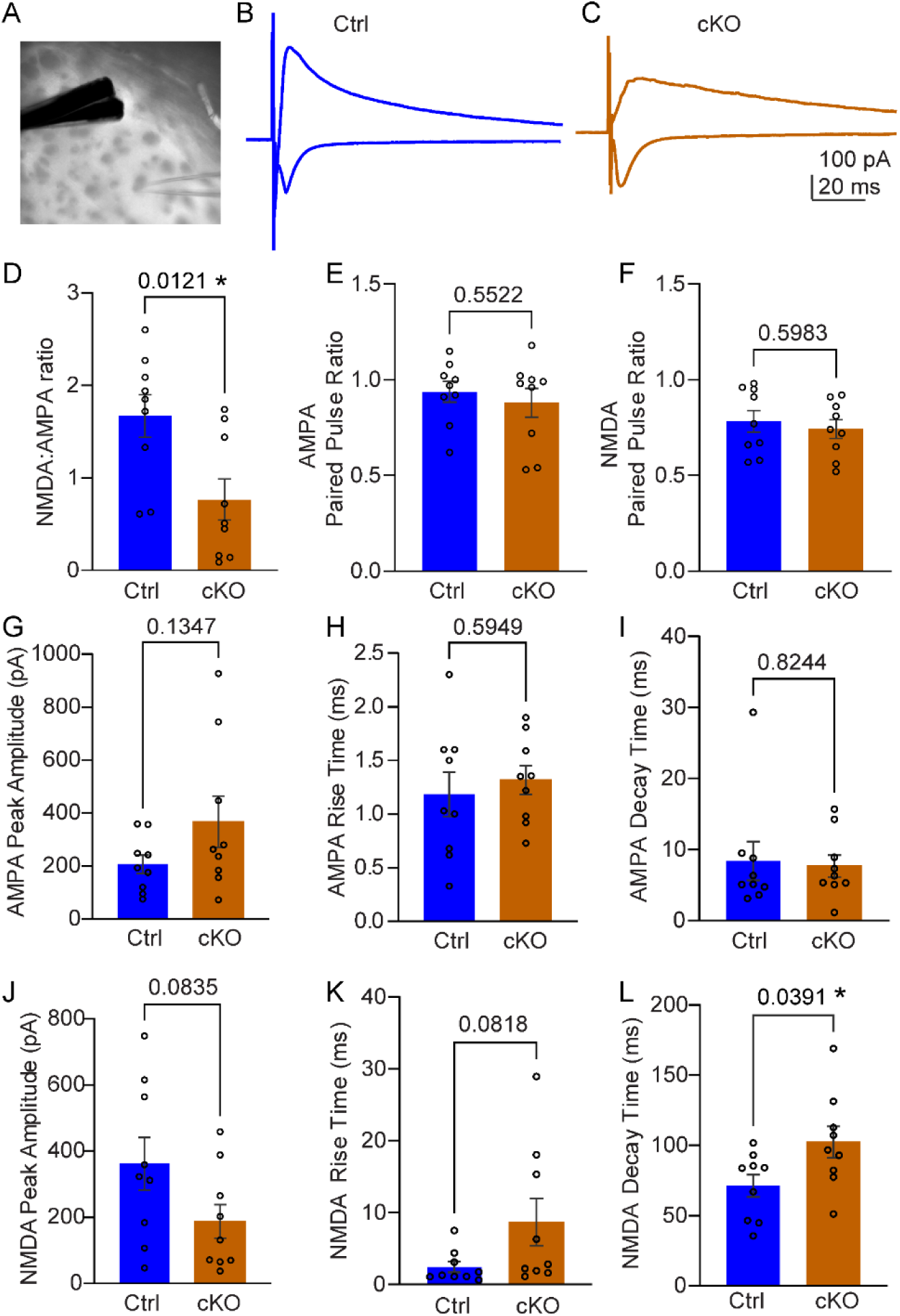
Cilia ablation reduces the NMDA:AMPA ratio and slows NMDA receptor decay in striatal MSNs. **(A)** Representative IR-DIC image of stimulating and whole cell patch clamp electrode placement. **(B)** Traces showing the average evoked AMPA receptor- and NMDA receptor-mediated currents recorded at −70 mV and +40 mV, respectively, from striatal neurons in Ctrl and **(C)** in cKO mice. **(D)** Bar graph showing NMDA:AMPA ratio. **(E)** AMPA current paired pulse ratio. **(F)** NMDA current paired pulse ratio. Data are presented as mean ± SEM with individual data points; Unpaired t-test. **(G)** Peak amplitude of AMPA receptor mediated responses. **(H)** Rise time of AMPA responses. **(I)** Decay time of AMPA responses. **(J)** Amplitude of NMDA currents. **(K)** Rise time of NMDA responses. **(L)** Decay time of NMDA responses. Data in bar graphs are presented as mean ± SEM with individual data points marked. n = 9 cells/4 mice per group, unpaired t-test p-values are displayed.

### Striatal cilia ablation produces region-specific metabolomic remodeling

Untargeted metabolomics profiling identified 714 annotated metabolites across STR and CTX from control and cKO mice, spanning nine super-pathways dominated by lipids (347 metabolites, 48.6%) and amino acids (161 metabolites, 22.5%) (Fig. 5A, Supplementary Dataset 1). Principal component analysis revealed that brain region was a major source of metabolic variance, with PC2 (15.93%) separating striatal from cortical samples regardless of genotype (Fig. 5B). One STR- IFT88-cKO sample with an extreme PC1 score was identified as an outlier and excluded from subsequent analyses. In control mice, the striatum and cortex already exhibited markedly different metabolomic profiles, with 303 of 714 metabolites (42.4%) significantly differing between regions in control animals. The striatum was substantially more specialized, with 251 metabolites significantly higher and 52 significantly lower than in the cortex (Fig. 5C,D). The most prominently striatum-enriched metabolites included dopamine (FC=38.8), 3-methoxytyramine (FC=28.2), and homovanillate (FC=6.6), consistent with the dopaminergic identity of the striatum. Lipids represented the largest class of regionally enriched metabolites (139 STR-enriched vs 29 CTX-enriched; Fig. 5D). Cilia ablation elicited markedly region-specific responses: 207 metabolites were altered in the striatum versus only 60 in the cortex (Fig. 5E,F, Supplementary dataset 1). The direction of change also differed strikingly between regions. In the striatum, 195 of 207 significantly altered metabolites (94.2%) were decreased, with only 12 increased. In contrast, the cortex showed the opposite pattern, with most changes being increases (47 increased versus 13 decreased; Fig. 5E,F). Comparison across regions identified 185 metabolites altered only in the striatum, 38 altered only in the cortex, and 22 shared (Fig. 5G,H). Together, these findings show that striatal cilia ablation produces a broad, predominantly downward, and regionally selective metabolomic phenotype.

**Figure 5.**
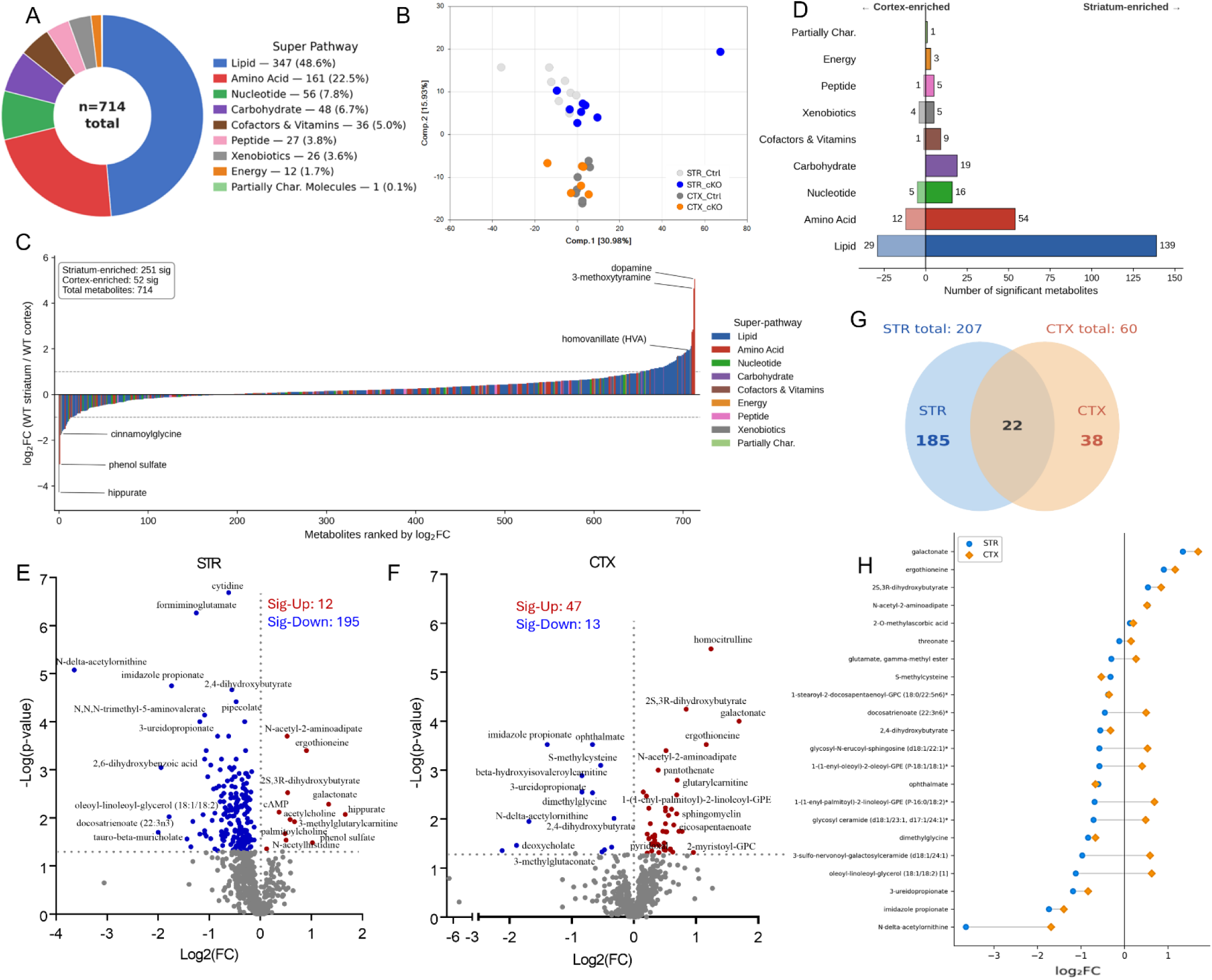
Striatal cilia ablation produces broad, regionally selective metabolomic depletion. **(A)** Donut chart showing the composition of the 714 annotated metabolites across nine super-pathways. **(B)** Principal component analysis (PCA) of all 714 metabolites across four groups (STR Ctrl, STR cKO, CTX Ctrl, CTX cKO). **(C)** Ranked dot plot of all 714 metabolites ranked by log₂FC (Ctrl STR / Ctrl CTX), illustrating the baseline metabolic specialization of the STR relative to CTX. Significantly enriched metabolites (p<0.05) are colored by super-pathway; non-significant metabolites are shown in grey. **(D)** Bar chart, showing the number of metabolites significantly altered in Ctrl STR vs Ctrl CTX, broken down by super-pathway. **(E, F)** Volcano plots showing significantly altered metabolites in cKO vs Ctrl in **(E)** STR and **(F)** CTX. **(G)** Venn diagram showing the overlap of significantly altered metabolites between STR and CTX. **(H)** Dot plot showing log₂FC (cKO/Ctrl) in STR and CTX for the 22 metabolites significantly altered in both regions. n = 8 Ctrl STR, n = 7 cKO STR, n = 7 Ctrl CTX, n = 7 cKO CTX.

### Lipid-associated pathways are prominently altered following striatal cilia ablation

Among the 207 significantly altered striatal metabolites, 117 (56.5%) belonged to the lipid super-pathway, followed by amino acids (44 metabolites, 21.3%) and nucleotides (16 metabolites, 7.7%; Fig. 6A,B; Supplementary dataset 1). When expressed as a proportion of all identified metabolites within each class, 33.7% of all measured lipid species were significantly altered in the striatum, compared with 27.3% of amino acids and 28.6% of nucleotides, indicating that the dominance of lipids reflects true pathway-level vulnerability rather than a consequence of their abundance in the measured metabolome. By contrast, the cortical response was smaller and more mixed across classes, with lipids still prominent but amino acids and cofactors/vitamins making larger relative contributions (Fig. 6C,D; Supplementary dataset 1).

**Figure 6.**
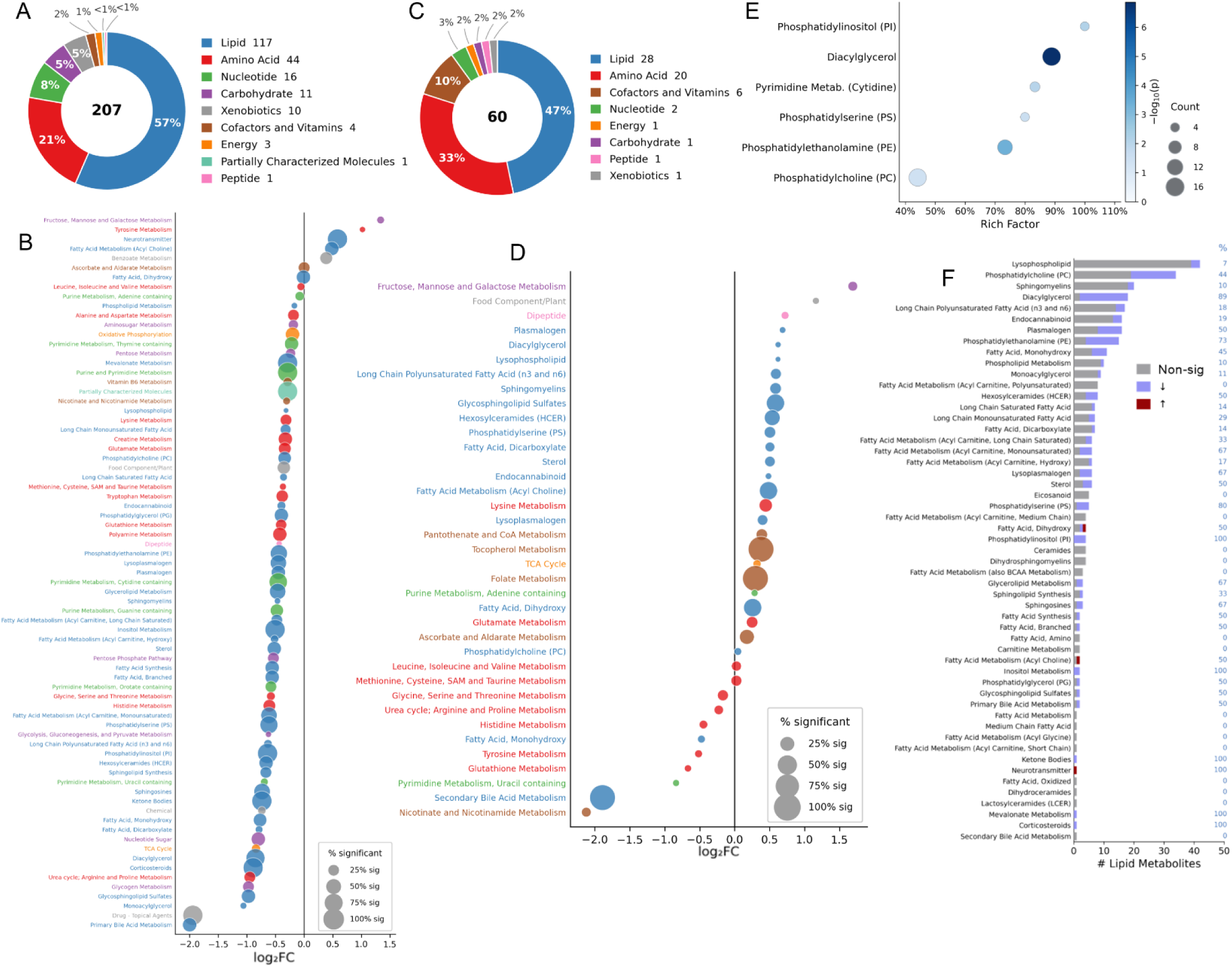
Striatal cilia ablation produces broad metabolomic remodeling dominated by lipid-associated metabolite reductions. **(A)** Donut chart showing the super-pathway distribution of 207 significantly altered metabolites in the striatum following cilia ablation. **(B)** Pathway enrichment dot plot for the STR, with pathways ranked by mean log₂FC (cKO/Ctrl). Dot size indicates the percentage of measured metabolites within each pathway that were significantly altered, and color indicates super-pathway class. **(C)** Donut chart showing the super-pathway distribution of significantly altered metabolites in cortex. **(D)** Pathway enrichment dot plot for the CTX, with pathways ranked by mean log₂ fold change (cKO/Ctrl). Dot size indicates the percentage of measured metabolites within each pathway that were significantly altered, and color indicates super- pathway class. **(E)** Pathway overrepresentation analysis (ORA) of significantly altered striatal metabolites, performed using a hypergeometric test. Rich factor represents the proportion of measured metabolites within each pathway that were significantly altered (significant/total metabolites per pathway); dot size indicates the number of significant metabolites, and dot color indicates −log₁₀(p). Significantly enriched pathways included diacylglycerol, phosphatidylethanolamine, phosphatidylinositol, cytidine-containing pyrimidine metabolism, phosphatidylserine, and phosphatidylcholine. **(F)** Bar plot showing the number of significantly decreased and increased metabolites across lipid sub-pathways in cKO STR relative to Ctrl striatum. Percentages on the right indicate the fraction of measured metabolites altered within each lipid sub-pathway.

To identify which sub-pathways were statistically overrepresented, we performed pathway overrepresentation analysis (ORA) using a hypergeometric test (Fig. 6E). Six sub-pathways were significantly overrepresented among the striatal hits (p<0.05), five of which were phospholipid classes: diacylglycerol (DAG; 16/18 species significantly decreased, RF=88.9%, p<0.0001), phosphatidylethanolamine (PE; 11/15, 73.3%, p=0.0003), phosphatidylinositol (PI; 4/4, 100%, p=0.006), phosphatidylserine (PS; 4/5, 80%, p=0.024), and phosphatidylcholine (PC; 15/34, 44.1%, p=0.031). The one non-lipid pathway was pyrimidine metabolism, cytidine-containing (5/6, 83.3%, p=0.008). Notably, all five phospholipid classes are major structural and signaling components of the cell membrane, and all significantly altered metabolites within each of these pathways were decreased without exception. This pattern of broad suppression extended across the majority of lipid sub-pathways, with decreased species predominating, while only a small subset showed increases (Fig. 6F). Together, these data indicate that the striatal metabolomic phenotype is dominated by broad and coordinated lipid-associated molecular remodeling, with membrane phospholipid pathways showing the most statistically robust enrichment.

### Striatal cilia ablation alters neurotransmitter-associated metabolites

Beyond the dominant lipid phenotype, prominent striatal changes involved neurotransmitter-associated metabolites, across several biologically interconnected pathways (Fig. 7A). The polyamine axis was broadly suppressed: spermine (FC=0.61; p=0.009), spermidine (FC=0.83; p=0.023), 4-acetamidobutanoate (FC=0.71; p=0.018), and 5-methylthioadenosine (FC=0.86; p=0.029) were all significantly decreased (4/8 pathway metabolites; Fig. 7A). Glutamatergic metabolites were coordinately reduced, with glutamate (FC=0.89; p=0.001), N-acetyl-aspartyl-glutamate (NAAG; FC=0.78; p=0.003), carboxyethyl-GABA (FC=0.70; p=0.001), and glutamate-γ-methyl ester (FC=0.81; p=0.005) all significantly decreased (4/10 pathway metabolites; Fig. 7A,B), along with ophthalmate (FC=0.66; p=0.001), a γ-glutamyl derivative and glutathione analog. The tryptophan-kynurenine pathway was similarly suppressed, with tryptophan (FC=0.82; p=0.006), indolelactate (FC=0.69; p=0.009), and kynurenine (FC=0.80; p=0.037) significantly reduced (3/8 pathway metabolites; Fig. 7A,C). Histidine metabolism was broadly affected (7/18 metabolites; Fig. 7A,D), with significant decreases in imidazole propionate (FC=0.30; p<0.001), formiminoglutamate (FC=0.42; p<0.001), histidine (FC=0.81; p=0.009), homocarnosine (FC=0.61; p=0.024), and 1-methylhistidine (FC=0.72; p=0.005), while N-acetylhistidine was significantly increased (FC=1.42; p=0.029). Purine metabolism was also altered, but in a divergent pattern (Fig. 7A,E). The purine nucleosides and bases were coordinately decreased — adenosine (FC=0.78; p=0.021), guanosine (FC=0.76; p=0.035), guanine (FC=0.71; p=0.002), and guanosine 5′-monophosphate (5′-GMP; FC=0.69; p=0.032), with guanosine 5′-diphosphate (GDP) trending lower (FC=0.82; p=0.074), whereas the downstream catabolites hypoxanthine, xanthine, and urate were unchanged, indicating reduced purine pools rather than enhanced terminal degradation. In contrast, the second messenger cyclic AMP was significantly increased (FC=1.29; p=0.008). A comparable increase was seen in the cholinergic transmitter acetylcholine (FC=1.50; p=0.011). Notably, dopaminergic metabolites were spared (tyrosine, dopamine, homovanillate, and 3- methoxytyramine), underscoring the selectivity of these changes.

**Figure 7.**
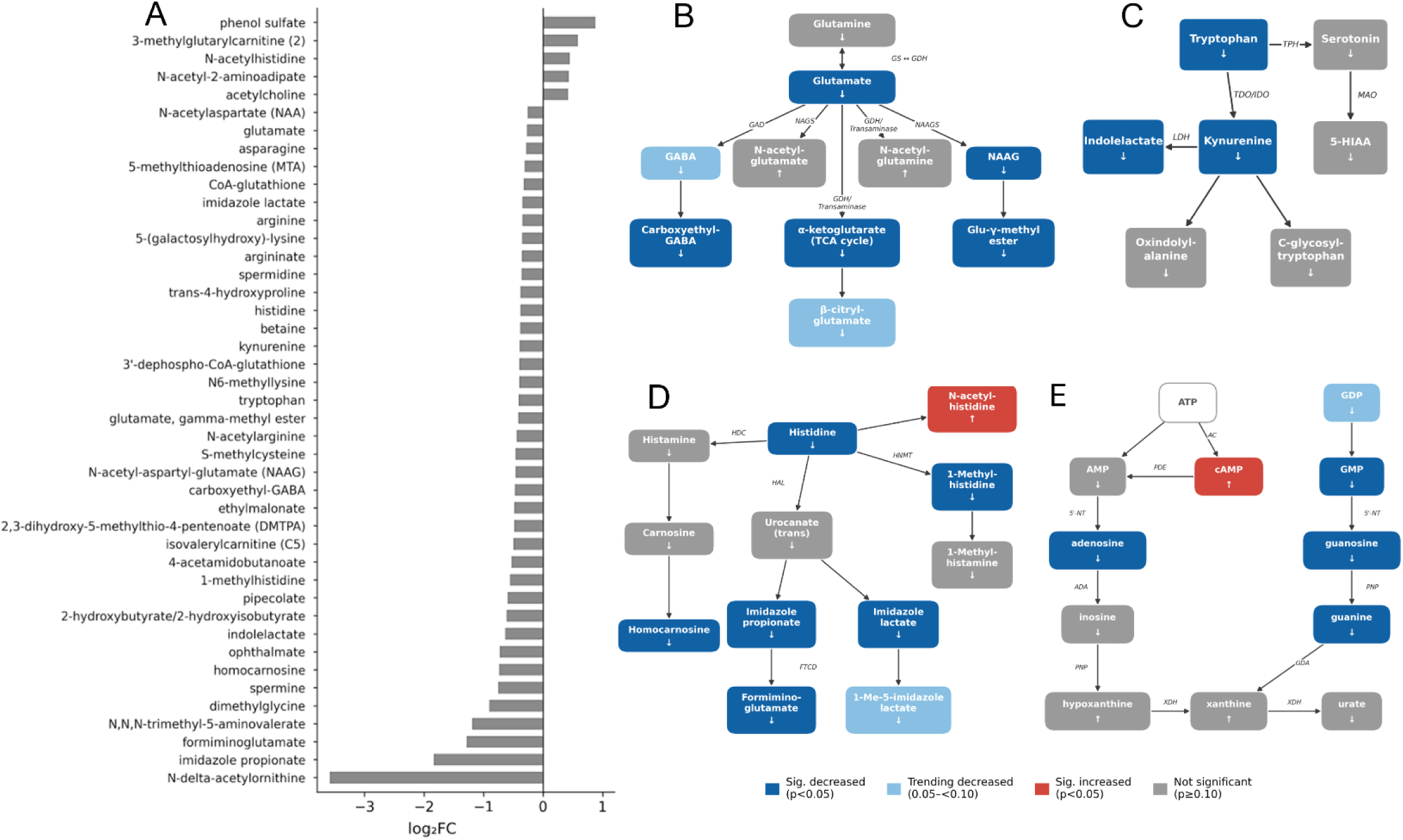
Striatal cilia ablation alters neurotransmitter-related metabolites and spontaneous activity patterning. **(A)** Bar plot showing fold-change values for neurotransmitter-related metabolites significantly altered in the striatum of cKO mice compared with Ctrl mice. Metabolites are ranked by effect size. **(B)** Glutamate-related pathway map showing altered metabolites in the striatum after cilia ablation. Boxes represent individual metabolites, and the arrow within each box indicates the direction of change. Box color denotes the direction and significance of change: dark blue, significantly decreased (p<0.05); light blue, trending decrease (0.05≤p<0.10); dark red, significantly increased (p<0.05); gray, not significantly changed (p≥0.10). Enzyme abbreviations are shown along pathway arrows: GAD, glutamate decarboxylase; NAGS, N-acetylglutamate synthase; GDH, glutamate dehydrogenase; GS, glutamine synthetase. **(C)** Tryptophan pathway map showing altered metabolites after striatal cilia ablation. Enzyme abbreviations: TPH, tryptophan hydroxylase; TDO/IDO, tryptophan 2,3-dioxygenase/indoleamine 2,3-dioxygenase; MAO, monoamine oxidase; LDH, lactate dehydrogenase. **(D)** Histidine pathway map showing altered metabolites after striatal cilia ablation. Enzyme abbreviations: HDC, histidine decarboxylase; HNMT, histamine N-methyltransferase; HAL, histidine ammonia-lyase; FTCD, formimidoyltransferase-cyclodeaminase. **(E)** Purine pathway map showing altered metabolites after striatal cilia ablation. ATP (open box) was not measured and is shown as the upstream precursor. Enzyme abbreviations: AC, adenylyl cyclase; PDE, phosphodiesterase; 5′-NT, 5′-nucleotidase; ADA, adenosine deaminase; PNP, purine nucleoside phosphorylase; GDA, guanine deaminase; XDH, xanthine dehydrogenase/oxidase.

### Striatal cilia ablation reduces activity at the dark-phase transition without altering overall locomotor activity

To determine whether striatal cilia loss affects the organization of spontaneous activity, home-cage locomotor activity was monitored across multiple light–dark cycles. Both Ctrl and cKO mice showed clear light– dark activity patterns, indicating preserved entrainment to the environmental cycle (Fig. 8A-C). cKO mice displayed reduced activity at selected time points, with the largest differences occurring around the transition into the dark phase. Activity was significantly reduced at hour 8 (Ctrl: 535.4 ± 32.4 vs. cKO: 277.3 ± 28.2 counts; p = 0.009) and hour 9 (Ctrl: 418.7 ± 45.2 vs. cKO: 240.6 ± 31.8 counts; p = 0.030; Fig. 7F-H), with additional reductions at hours 5, 11, and 24. These results suggest that striatal cilia ablation particularly affects the normal surge in activity at the dark-phase transition.

**Figure 8.**
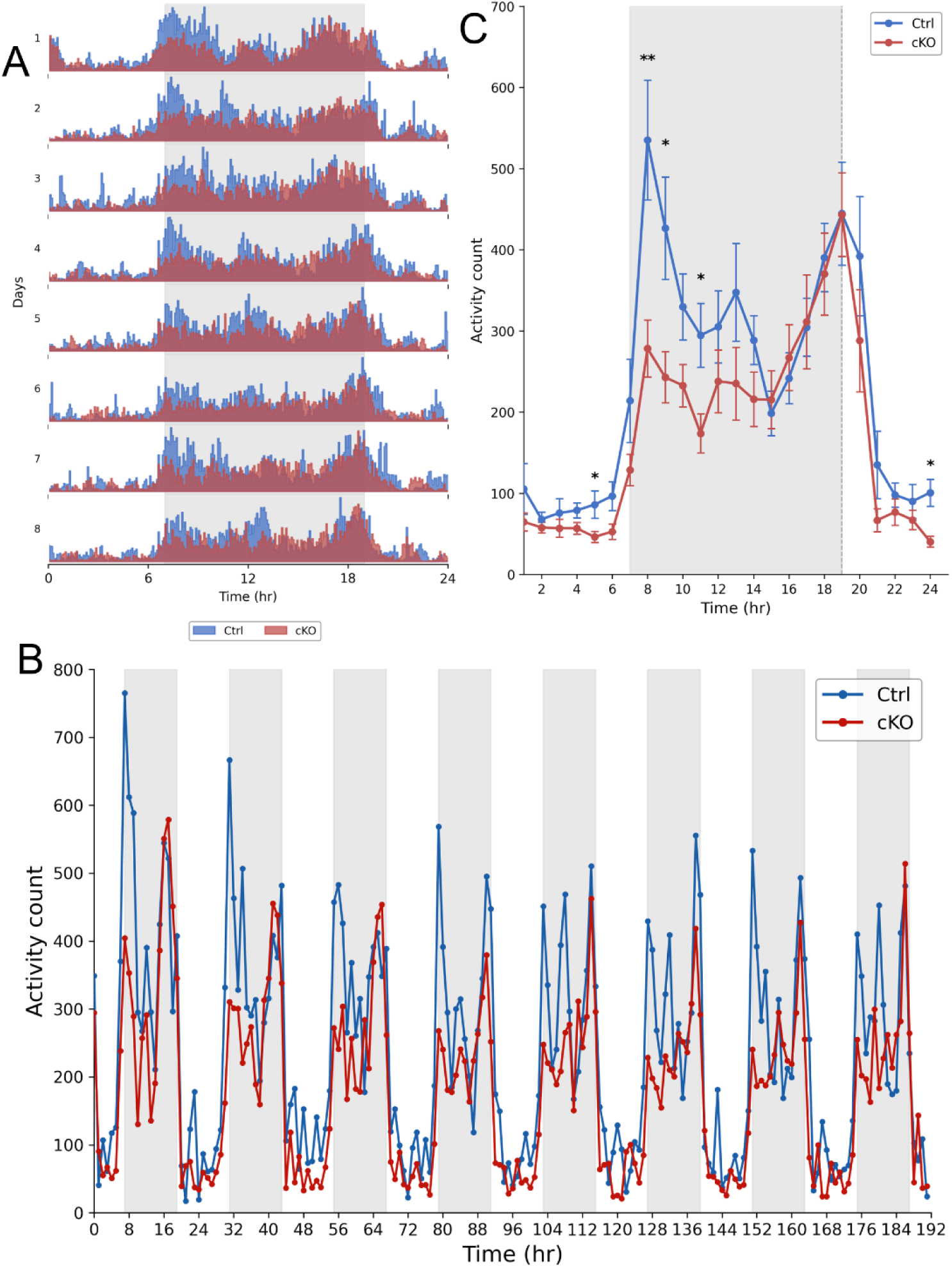
Striatal cilia ablation alters spontaneous activity patterning, predominantly during light-dark transition. Spontaneous locomotor activity in Ctrl and Ift88-cKO mice recorded over 8 consecutive days using an infrared photobeam system. **(A)** Double-plotted actogram showing activity counts across repeated 24-h light–dark cycles. **(B)** Continuous activity counts across the full 8-day recording period, plotted in 1-h bins. **(C)** Mean hourly activity profile averaged across recording days. x-axis indicates clock time; gray shading indicates dark phase, 07:00–19:00. Two-way ANOVA: genotype effect, F(1,312) = 29.18, p<0.0001; time-of-day effect, F(23,312) = 22.49, p<0.0001; followed by post hoc comparisons; *p<0.05, **p<0.01. Data are presented as mean ± SEM; Ctrl n=8, cKO n=7.

## DISCUSSION

In this study, we examined the role of the primary cilium in striatal neuron connectivity and function. By combining selective *Ift88* deletion with monosynaptic rabies tracing, whole-cell electrophysiology, and untargeted metabolomics, we show that cilia loss produces a 3-fold reduction in monosynaptic afferent convergence, weakened excitatory synaptic transmission at surviving synapses, and broad, region-selective metabolomic remodeling. Together, these findings identify primary cilia as regulators of the biochemical environment that supports striatal excitatory synaptic functions and connectivity.

### Primary cilia shape the architecture of striatal afferent connectivity

A striking anatomical finding was the scale and selectivity of afferent loss following striatal cilia ablation. Total monosynaptic input to cilia-deficient MSNs was reduced approximately threefold, yet this reduction was not distributed uniformly across brain projection systems. Cognitive-associative and somatosensory inputs showed large reductions, whereas motor execution inputs were comparatively spared. Basal ganglia inputs were also significantly reduced, but to a lesser extent than cortical and thalamic inputs. This pattern of selective vulnerability aligns with the behavioral profile we previously reported in the same mouse line, in which striatal cilia deletion impaired working memory, sensorimotor gating, motor learning, and repetitive behavior, while sparing spontaneous locomotion [23]. Consistent with this dissociation, cKO mice maintained overall locomotor activity but showed a reduced activity response at the transition to dark-phase. Thus, the behavioral effect of striatal cilia loss appears strongest when animals must quickly adjust activity to changing conditions, rather than during basal movement. The impaired behaviors likely depend on integration of associative cortical, somatosensory, and higher-order thalamic signals, the afferent domains most affected by cilia loss, whereas relatively preserved motor execution pathways may explain the sparing of basic locomotor output [1–4]. The tracing data may, therefore, provide a circuit-level framework for a behavioral dissociation that previously lacked an anatomical basis.

### Surviving synapses are functionally weakened after cilia loss

Whole-cell recordings revealed that the anatomical loss of afferent inputs is accompanied by functional weakening at the remaining synapses. Reduced mEPSC frequency is consistent with fewer functional excitatory contacts, in agreement with the monosynaptic tracing data. Although reduced mEPSC amplitude can be hard to interpret, it is consistent with impaired postsynaptic excitatory transmission at surviving synapses[26]. Evoked synaptic recordings further showed a reduced NMDA:AMPA ratio and slower NMDA receptor kinetics in cKO mice, indicating a selective alteration in NMDA receptor-mediated transmission. One possible interpretation is that cilia loss reduces synaptic NMDA receptor contribution and may alter NMDA receptor subunit composition, potentially increasing the relative contribution of slower GluN2B-containing receptors [27–29]. Together, these findings demonstrate that loss of primary cilia weakens excitatory synaptic transmission at multiple levels, with fewer remaining functional inputs showing diminished NMDA receptor signaling.

Importantly, these electrophysiological results extend the anatomical findings from monosynaptic tracing. While rabies tracing reveals the architecture of afferent connectivity, it cannot assess whether the remaining synapses are functionally competent. Our recordings show that the surviving synaptic network is not only reduced in size but is also functionally weakened. This indicates that primary cilia contribute to the functional maintenance of excitatory synapses, rather than solely supporting anatomical connectivity.

### Metabolomics links cilia loss to synaptic and circuit remodeling

Metabolomic profiling revealed pronounced and regionally selective changes following striatal cilia ablation. The striatum exhibited widespread metabolite reductions, whereas the cortex showed fewer and often directionally opposite changes. This regional asymmetry in metabolomic changes indicates that the metabolic effects of cilia loss were largely confined to the targeted striatum rather than reflecting a nonspecific, brain-wide metabolic disturbance.

Several altered biochemical pathways map directly onto the observed synaptic phenotype. First, cilia ablation produced a broad lipid-associated signature, marked by reductions in major membrane lipid classes, including DAG, phosphatidylcholines, and phosphatidylethanolamines. DAG was the most significantly overrepresented pathway, with 16 of 18 detected species decreased. Beyond its role as a structural membrane component, DAG is also a principal activator of Protein Kinase C (PKC) [30], which phosphorylates GluA1 at Ser831 to drive AMPA receptor insertion at the postsynaptic density [31, 32]. Therefore, depletion of the DAG pool could, probably, impair AMPA receptor trafficking and contribute to reduced mEPSC amplitude. This provides a plausible molecular link between lipid-associated metabolic disruption and weakened postsynaptic excitatory transmission. Polyamine metabolism provides a second link. Spermidine and spermine were significantly reduced, tracing upstream to near-complete loss of N-delta-acetylornithine, the most strongly depleted metabolite in the dataset. Polyamines can potentiate GluN2B-containing NMDA receptors by enhancing channel open probability without necessarily altering receptor kinetics [33, 34]. Their depletion could therefore contribute to the reduced NMDA receptor-mediated current and NMDA:AMPA ratio. Because polyamine modulation acts on channel open probability rather than gating kinetics, however, the slowed NMDA decay we observed likely reflects an additional mechanism, such as a shift in subunit composition toward slower GluN2B-containing receptors.

A third link involves glutamate-associated metabolites. Glutamate, NAAG, and NAA were all reduced in the striatum. Altered glutamate availability may directly affect excitatory transmission [35, 36], while changes in NAAG and NAA point to disrupted glutamate handling and axonal metabolic support [37, 38]. These changes could contribute to reduced mEPSC frequency and may also relate to the anatomical reduction in afferent input. Thus, the DAG-PKC-GluA1 axis, polyamine- NMDA receptor modulation, and glutamate/NAAG/NAA metabolism each provide plausible bridges between the metabolomic signature and specific electrophysiological observations. Together, these findings suggest that cilia loss disrupts the molecular infrastructure required for synaptic maintenance. Rather than representing an isolated metabolic phenotype, the metabolomic signature provides a biochemical context for the observed weakening of synaptic transmission and contraction of afferent architecture. Notably, these reductions occurred alongside selective increases in cAMP and acetylcholine, suggesting that cilia loss produces a broader neurochemical remodeling of the striatum, marked by reduced glutamate-related metabolites but increased cholinergic and cyclic nucleotide signaling. The elevation of cAMP is interesting given that AC3, the primary adenylyl cyclase isoform in neuronal cilia, is lost with cilia ablation, a finding that points to complex rebalancing of cAMP signaling between ciliary and non-ciliary compartments rather than a simple reduction in cAMP production. The increase in acetylcholine may reflect altered striatal cholinergic tone, potentially arising from disrupted MSN–cholinergic interneuron interactions or compensatory network adaptation following reduced excitatory signaling.

### Primary cilia as biochemical regulators of striatal circuit integrity

The ciliary membrane concentrates a specialized GPCR repertoire, including Drd1, Drd2, 5-HT6, Mc4r, and Gpr88 [10, 14, 16–19, 39, 40], whose signaling is spatially organized within the ciliary compartment and distinct from somatic receptor pools. Loss of this compartment may, therefore, disrupt the spatial organization of GPCR signaling rather than simply eliminating it. several cilia-enriched receptors can also signal from non-ciliary membrane domains, and their redistribution following cilia loss may shift the balance of downstream signaling cascades in ways that are not yet fully characterized. The observed reductions in DAG and phospholipid-related metabolites, alongside increases in cAMP and acetylcholine, are consistent with such a rebalancing, though whether these changes reflect direct consequences of ciliary GPCR loss or secondary network adaptations cannot be determined from the present data.

The convergence of anatomical, electrophysiological, and metabolomic findings points to a linked disruption in which the biochemical state of MSNs influences their capacity to maintain excitatory synapses and preserve afferent convergence. Whether the metabolomic changes are causally upstream of the synaptic and anatomical deficits, or whether all three reflect parallel consequences of cilia loss, cannot be determined from the present data. Reduced DAG could impair PKC-dependent AMPA receptor trafficking; reduced polyamines could decrease NMDA receptor potentiation; and reductions in glutamate and NAAG could limit excitatory transmission and presynaptic release. These relationships suggest that the metabolomic signature is functionally relevant to the synaptic phenotype, rather than a separate consequence of cilia loss. Together, these findings position the primary cilium as a regulator of the molecular environment that supports excitatory synaptic transmission and connectivity in striatal MSNs.

### Implications for circuit dysfunction in neurological and psychiatric diseases

Disrupted corticostriatal and thalamostriatal connectivity is a prominent feature of several neurological and psychiatric disorders, including Parkinson’s disease [41, 42], Huntington’s disease [43, 44], schizophrenia [3, 45], obsessive-compulsive disorder [46], addiction [47], and autism spectrum disorder [3, 45]. Circuit dysfunction is also relevant to ciliopathies such as Joubert syndrome and Bardet-Biedl syndrome, which present with cognitive and motor phenotypes, whose neural basis remains incompletely understood [48–50]. The present findings suggest that ciliary dysfunction may contribute to circuit pathology not only through altered development, but also through impaired maintenance of adult neural connectivity.

This distinction is important. Primary cilia have traditionally been viewed through the lens of developmental signaling, however, the present study shows that their loss in adult striatal neurons is associated with metabolic remodeling, weakened excitatory synaptic function, and selective contraction of brain-wide afferent input. These findings support a model in which cilia continue to regulate circuit integrity after development by maintaining the molecular environment required for synaptic and anatomical stability.

More broadly, this study raises the possibility that ciliary dysfunction may represent an underappreciated mechanism of progressive circuit deterioration in disorders marked by loss of associative corticostriatal and thalamostriatal connectivity [9, 11]. Whether restoration of ciliary signaling in adult animals can reverse the metabolic, synaptic, and anatomical consequences of cilia loss remains an important open question with direct therapeutic relevance.

## MATERIALS AND METHODS

### Animals

Ift88*fl/fl* mice carrying loxP sites flanking exons 4–6 of the *Ift88* gene (Jackson Laboratories, stock #022409) were maintained on a C57BL/6J background and bred as Ift88^fl/+^ × Ift88^fl/+^ intercrosses. Homozygous Ift88^fl/fl^ mice and wild-type (Ift88^+/+^) littermates, which served as controls, were used throughout the experiments and received identical viral injections. Genotypes were confirmed by PCR. Only male mice were included. All animal procedures were approved by the IACUC at the University of California, Irvine, and conducted in accordance with institutional and national ethical guidelines.

### Monosynaptic Rabies Virus Tracing

Ift88*fl/fl* and control mice received stereotaxic co-injection of AAV9-CaMKII0.4-Cre-SV40 (0.2 μL; 1.3×10¹³ GC/mL; Addgene #105558) and a 1:1 mixture of AAV5-CAG-FLEX-TC66T- mCherry and AAV8-CAG-FLEX-RABV-G (0.3 μL total; 1.9×10¹² GC/mL each) into the dorsal striatum (+1.3 mm AP, +1.5 mm ML, −3.0 mm DV from bregma). Fourteen days later, 0.5 μL of EnvA-pseudotyped RABVΔG-GFP was injected at the same coordinates. Brains were collected 5 days after rabies injection following transcardial perfusion with 4% PFA. Coronal sections (30 μm) were immunostained with anti-ADCY3 (1:500; cilia marker) and anti-mCherry (1:500) and imaged at 4× on an Olympus IX83 and 20× on a Zeiss LSM 900 confocal. Starter cells were TVA-mCherry⁺/GFP⁺ double-positive neurons; input neurons were GFP⁺/DAPI⁺/mCherry⁻. Every sixth section, corresponding to approximately 180 μm spacing, was sampled. Anatomical assignments followed the Allen Mouse Brain Atlas. Input strength was expressed as STSta (%) = (GFP⁺ input cells / starter cells) × 100. Global input convergence was compared by unpaired two-tailed t-test; regional comparisons used unpaired two-tailed t-tests. All analyses were performed in GraphPad Prism 10.4. n = 3/group.

### Whole-Cell Patch-Clamp Electrophysiology

Seven to ten days post-surgery, control or Ift88-cKO mice were briefly anesthetized with isoflurane and transcardially perfused with warm (30–32°C) cutting solution containing (in mM): 110 choline, 25 NaHCO3, 1.25 NaH2PO4, 3 KCl, 7 MgCl2, 0.5 CaCl2, 10 glucose, 11.6 sodium ascorbate, and 3.1 sodium pyruvate, bubbled with 95% O2 and 5% CO2. 400 μm thick slices containing the dorsal striatum, specifically to include the injected sites, were prepared on a Z-deflection calibrated vibrating blade microtome (Campden SMZ 7000-2, Lafayette Instruments). Slices were maintained in the cutting solution for an additional 15 minutes at 32°C. After incubation, slices were transferred to artificial cerebrospinal fluid (ACSF) containing (in mM): 126 NaCl, 26 NaHCO3, 1.25 NaH2PO4, 3 KCl, 1 MgCl2, 2 CaCl2, 10 glucose, bubbled with 95% O2 and 5% CO2, and kept in a holding chamber at room temperature for 20–30 minutes before use [51, 52].

All experiments were performed at near physiological temperature (32–34°C) in a submersion-type recording chamber mounted on an Olympus BX51-WI microscope. Whole-cell patch-clamp recordings were obtained from medium spiny neurons in dorsal striatum slices with positive GFP expression, indicating transduction of AAV9.CamKII.HI.GFP- Cre.WPRE.SV40 in either wild-type control or Ift88^fl/fl mice.

Miniature excitatory postsynaptic currents (mEPSCs) were recorded in voltage clamp mode with bath application of 1 μM Tetrodotoxin (TTX, Tocris) and 50 μM Picrotoxin (PTX, Tocris) to block action potential-mediated synaptic currents and GABAAR-mediated currents, respectively. Glass electrodes (2–4 MΩ) were filled with internal solution containing (in mM): 135 KMeSO3, 10 HEPES, 4 MgCl2, 4 Na2ATP, 0.4 NaGTP, and 10 sodium creatine phosphate, pH adjusted to 7.3 with KOH.

Evoked excitatory postsynaptic currents (eEPSCs) were elicited by stimulating local afferents using a bipolar stimulating electrode placed in the dorsal striatum approximately 100–150 μm from the recorded MSN. Stimulation intensity was adjusted to evoke reliable postsynaptic responses (typically 250–300 μA, 500 μs duration). AMPA receptor-mediated currents were recorded at a holding potential of -70 mV. NMDA receptor-mediated currents were isolated by holding cells at +40 mV. Inhibitory currents through GABAa and GABAb receptors were blocked (PTX, 50 μM, and CGP55845, 1 μM) to pharmacologically isolate excitatory responses. For recording evoked AMPA and NMDA currents, internal solution with cesium was used to improve space clamp (in mM): 135 CsMeSO3, 10 HEPES, 4 MgCl2, 4 Na2ATP, 0.4 NaGTP, and 10 sodium creatine phosphate, pH adjusted to 7.3 with CsOH. Series resistance ranged from 10 to 25 MΩ.

Electrophysiological data were acquired using a MultiClamp 700B Amplifier (Molecular Devices), filtered at 4 kHz, and digitized at 10 kHz. Series resistance was monitored on all sweeps by evoking a brief 10mV pulse, and the data from analysis were discarded if this series resistance was greater than 20% from the beginning of the recording. Data collection was performed with National Instruments DAQ boards and Wavesurfer software written in MATLAB (Howard Hughes Medical Institute Janelia Research Campus). Offline analysis was done using Clampfit 10.7.

*Analysis and Statistics:* The experimenter was blinded to cKO condition during physiology experiments until data analysis was finished, then unblinded for statistical analysis. Individual mEPSC events were detected using a template-based search function in Clampfit (Molecular Devices). A representative event waveform was first selected and used to generate a template by averaging several manually identified events of similar shape and amplitude. The template was then applied to the entire recording using the template search function, which detects events based on the degree of correlation between the data and the template waveform. Events were reviewed visually, and any noise that falsely met trigger criteria was rejected. Cumulative probability plots of mEPSC parameters were generated by pooling an equal number of mEPSCs (35 events) from each cell from both the control and Ift88-cKO groups. Peak amplitudes of AMPA and NMDA currents were measured by averaging 5-10 sweeps per cell. From these averages, the maximum AMPA current peak amplitude was measured from the baseline, while the NMDA peak amplitude was measured ∼50ms after the stimulus onset or the AMPA peak. The NMDA to AMPA ratio was calculated using these peak amplitude values. Kinetics of AMPA and NMDA currents were analyzed using the statistics module in Clampfit software. Currents were peak-scaled and rise time was defined as the interval from 20% to 80% of the peak amplitude, while decay time as the interval from 80% to 20% of the peak.

Statistical analysis was conducted using paired and unpaired Student’s t-test or the Kolmogorov–Smirnov (K–S) test, depending on the data distribution. Significance was set at p < 0.05. Data are presented as mean ± SEM or median with interquartile range (IQR) where appropriate. All the statistical analyses and graphs were generated using GraphPad Prism 10.4.1.

### Home Cage Locomotor Activity Recording

Home-cage locomotor activity was recorded in Ctrl and cKO mice using standard home cages placed within locomotor activity chambers (40 × 40 × 38 cm³) equipped with a 16 × 16 photobeam arrays (San Diego Instruments, San Diego, CA). Mice were individually housed and allowed to acclimate to the recording chambers for 48 h before data collection. Activity was then recorded continuously in 5-min bins for 8 days under a 12:12 h light–dark cycle, with lights off at 07:00, lights on at 19:00. After activity recording, mice were returned to standard 12:12 h light–dark housing for two weeks before tissue collection for untargeted metabolomics. Activity counts were averaged across the 8 recording days and binned into 1-h intervals. Hourly activity scores were compared between groups using two-way ANOVA. Sample sizes were Ctrl n = 8 and Ift88-cKO n = 7.

### Untargeted Metabolomics

Brain tissue samples from striatum and cortex were collected from Ctrl and cKO mice and submitted to Metabolon for untargeted metabolomics profiling. Upon receipt, samples were accessioned into the Metabolon Laboratory Information Management System and stored at −80°C until processing. Samples were prepared using the automated MicroLab STAR system. Recovery standards were added before extraction for quality-control purposes. Proteins were precipitated with methanol under vigorous shaking, followed by centrifugation, to remove protein and recover chemically diverse small molecules. The resulting extracts were divided into five fractions: two for reverse-phase UPLC-MS/MS analysis in positive electrospray ionization mode, one for reverse-phase UPLC-MS/MS analysis in negative electrospray ionization mode, one for HILIC/UPLC-MS/MS analysis in negative electrospray ionization mode, and one backup fraction. Extracts were dried to remove organic solvent and stored under nitrogen before analysis.

Metabolites were analyzed by ultrahigh-performance liquid chromatography–tandem mass spectrometry using a Waters ACQUITY UPLC coupled to a Thermo Scientific Q-Exactive high-resolution accurate-mass spectrometer equipped with a heated electrospray ionization source and Orbitrap mass analyzer. Separate chromatographic conditions were used to optimize detection of hydrophilic and hydrophobic compounds, including acidic positive-ion reverse-phase methods, a basic negative-ion reverse-phase method, and a negative-ion HILIC method. The mass spectrometer alternated between MS and data-dependent MS/MS scans, with a scan range covering approximately 70–1000 m/z.

Raw data were extracted, peak-identified, and quality-control processed using Metabolon’s informatics pipeline. Metabolites were identified by comparison with library entries of purified standards based on retention index, accurate mass match within ±10 ppm, and MS/MS spectral matching. Peak areas were used for relative quantification. Quality control included pooled matrix samples, process blanks, solvent blanks, recovery standards, and internal standards. Experimental samples were randomized across the platform run, with QC samples spaced throughout the run to monitor instrument and process variability.

For statistical analysis, metabolite abundance values were log-transformed before comparison. Principal component analysis was used to assess global variation across samples. Differential metabolite abundance was analyzed by brain region and genotype. Pathway overrepresentation analysis was performed on Metabolon sub-pathway assignments using a one-sided hypergeometric test (scipy.stats.hypergeom; SciPy v1.17.1, Python v3.12.3) [53], with scripting refining assistance from Claude (Anthropic). Enrichment p-values were corrected for multiple comparisons using the Benjamini-Hochberg method. Rich factor is defined as the proportion of measured metabolites in a sub-pathway that were significantly altered. Only brain samples from striatum and cortex were included in the analyses reported here.

### General Statistical Analysis

All data are presented as mean ± SEM. Significance threshold: p < 0.05 (* p < 0.05, ** p < 0.01, *** p < 0.001). Experiment-specific statistical tests are described within each subsection above.

## Acknowledgements and funding sources

The work of AP and GL was supported by 1R01NS127785.

## Author contributions

KC designed and performed monosynaptic tracing experiments. KB designed and supervised monosynaptic tracing experiments. XF, RVM, and JL performed imaging and monosynaptic data analysis. AP and GL designed, performed, and analyzed electrophysiological experiments. SA and WA designed and performed motor activity and prepared brain tissues for metabolomic experiments. DS and SN contributed to metabolomics analysis. TD and SN contributed to manuscript data analysis and manuscript writing. AA conceived the study, designed the experiments, and supervised the project.

## AI Use statement

The authors used Claude (Anthropic) as an AI-assisted tool to support manuscript editing, code refinement, and language editing. Schematic elements in Figure 1 were partially generated using ChatGPT (OpenAI). All AI-generated content was reviewed, verified, and edited by the authors, who take full responsibility for the accuracy and integrity of the published work.

## Data availability

All data supporting the findings of this study are available within the article and its supplementary materials, and supplementary datasets.

